# bRaw: an R package for digital raw canopy imagery

**DOI:** 10.1101/2022.10.25.513518

**Authors:** F. Chianucci

## Abstract

Digital photography is an increasingly popular tool to estimate forest canopy attributes. However, estimates of gap fraction, upon which calculations of canopy attributes are based, are sensitive to photographic exposure in upward-facing images. Recent studies have indicated that analyzing RAW imagery, rather than other inbuilt camera format (e.g. jpeg, png, tiff) allows to obtain largely-insensitive gap fraction retrieval from digital photography. The package bRaw implemented the method proposed by Macfarlane et al. (2014). They found that shooting raw with one stop of underexposure and applying a linear contrast stretch yielded largely insensitive results, thus providing a way for standardizing and optimizing photographic exposure. The package replicate the methodology and thus it provides an effective tool to use raw imagery in canopy photography.

## Introduction

Digital photography is an increasingly popular tool to estimate forest canopy attributes. For simplicity, canopy photographic methods can be grouped into those based on wide-angle, multiple views and those based on a single, restricted-view. Wide-angle methods include circular fisheye (also called hemispherical) photography, which has been the pioneering method in canopy photography, with the first applications dating back about seventy years ago (Anderson (1964); Evans and Coombe (1959)). Full-frame fisheye photography also employed a fisheye lens, but it has lower field of view (FOV) compared to circular one, such that the full zenith angle range extends to the corners of the rectangular image, increasing the image resolution (Macfarlane et al. 2007b). Fifty-seven degree (57°) photography samples a restricted portion of the canopy centered at 1 radian, since canopy attributes at this view are not influenced by leaf orientation (Warren Wilson 1960). Finally, cover photography is a zenith observation method (Macfarlane et al. (2007a)), which is based on photographs acquired using a restricted 30° FOV. For a review on these methods, see Chianucci (2019).

Notwithstanding the wide use of digital canopy photography, estimates of gap fraction, upon which calculation of canopy attributes are derived, are sensitive to camera exposure in upward-facing images. A previous study (Beckschäfer et al. (2013)) indicated that at least ten different methods to determine exposure for canopy photography were used by scientists in the last two decades, hindering comparability among different studies and protocols.

**Figure 1.**
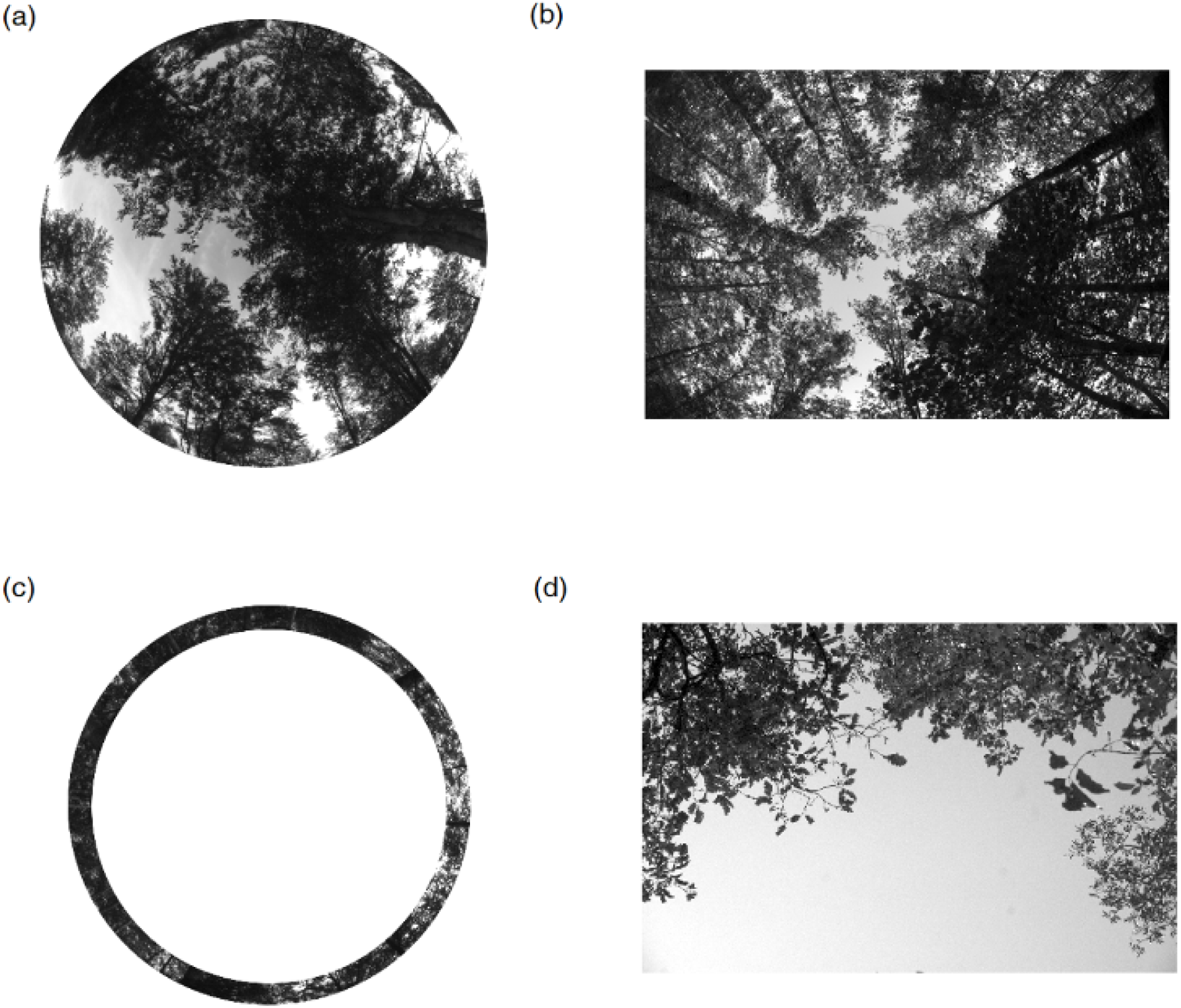
Example of different canopy photographic techniques: circular fisheye (a), full-frame fisheye (b), fifty-seven degree (c) and cover (d) photography. For description of the techniques, see Chianucci (2019)

Rather than looking for an optimal exposure from in-camera JPEG images, shooting raw has the advantage of higher radiometric resolution (bit-depth ≥ 12 bit) and linear relationship with actual brightness. Macfarlane et al. (2014) found that shooting raw with one stop of underexposure and applying a linear contrast stretch yielded largely insensitive results, thus providing a way for standardizing and optimizing photographic exposure.

The package `bRaw` implemented the method proposed by Macfarlane et al. (2014). The next paragraphs describe in detail the methodology and the package functioning.

### Brief description of the methodology

The bRaw package uses the functionality of dcraw, a program developed by Dave Coffin (available at: https://www.dechifro.org/dcraw/). Using the tool, linear readings of pixels are extracted from raw output files of the cameras; no interpolation (demosaiced) is applied, and the files are saved as simple portable gray map (PGM) files. This is equivalent to setting the following arguments in dcraw:

- −4 (write a linear 16 bit file);
- -D (document mode without interpolation nor scaling);
- -t 0 (no rotation).

By reading the Bayer pattern (-v -i), only the blue channel is then considered for subsequent analysis, as the blue portion of the spectrum allows best contrast between canopy and sky (Macfarlane et al. (2014)).

A linear contrast stretch is then applied to the non-interpolated subset of blue pixels, by saturating 1% of pixels at either ends of the blue histogram:

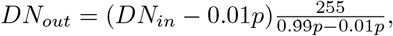

where *DN_in_* is the input pixel value, *DN_out_* is the output pixel value, 0.99*p* and 0.01*p* are the 99% and 1% percentile, respectively.

In case of circular fisheye image, the contrast stretch was based on the brightness of pixels within the circular mask only.

An optional gamma adjustment to 2.2 can also be applied to linear data:

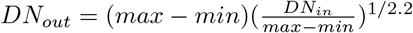

where *max* and *min* are the maximum and minimum values, respectively.

Finally, the blue channel is exported as 8-bit JPEG image; this results in a ‘lighter’ image, as only the blue fraction of the pixels (which typically corresponds to one-fourth of the native raw image resolution) are exported from original raw imagery.

### Installation

The `bRaw` package uses the functionality of dcraw, a raw-developing tool developed by David Coffin.

For Macintosh operating system, dcraw can be installed using the following command from the Terminal: `brew install dcraw`

For Windows operating system, ‘dcraw.exe’ must be downloaded from here and the file must be moved to the following location: `C:\DCRaw\`. The installer must be named ‘dcraw.exe’. The path to the image files must NOT contain spaces.

Once installed, it is possible to install `bRaw` using devtools (Wickham et al. 2021):

~~~
# install.packages(“devtools”)
devtools::install_gitlab(“fchianucci/bRaw”)
~~~

### Package usage

The key steps of the procedure proposed by Macfarlane et al. (2014) are:

1. read the Bayer pattern from RAW imagery;
2. convert the raw image into a 16 bit portable grey map (‘pgm’) format;
3. select the blue channel of the pgm;
4. (optionally) apply a circular mask (in case of circular fisheye images);
5. contrast stretch the image (or mask);
6. (optionally) apply a gamma adjustment (2.2);
7. Export the 16-bit linear, enhanced blue channel as a 8-bit single channel ‘jpeg’.

The `raw_blue` functions allows to perform all the processing steps above:

~~~
raw_blue(filename,circ.mask=NULL,gamma.adj=TRUE,display=TRUE)
~~~

where:

- *filename* is the input raw image;
- *circ.mask* is an optional argument, and correspond to the three-parameters (xc,yc,rc) required to set a circular mask (in case of circular fisheye images);
- *gamma.adj* is an optional argument, which allows to apply a gamma adjustement prior to exporting jpeg image;
- *display* allows the user to plot a graph of the image, along with the applied mask.

The function creates a ‘bRaw’ folder, where the exported 8-bit image is stored.

### Example usage

We illustrate the use of the function with a full-frame fisheye RAW image.

~~~
library(bRaw)
filename<-system.file(‘extdata/531539_AGO12_531.NEF’,package=‘bRaw’)
#> [1] “/Users/francescochianucci/Library/R/x86_64/4.1/library/bRaw/extdata/531539_AGO12_531.pgm”
~~~

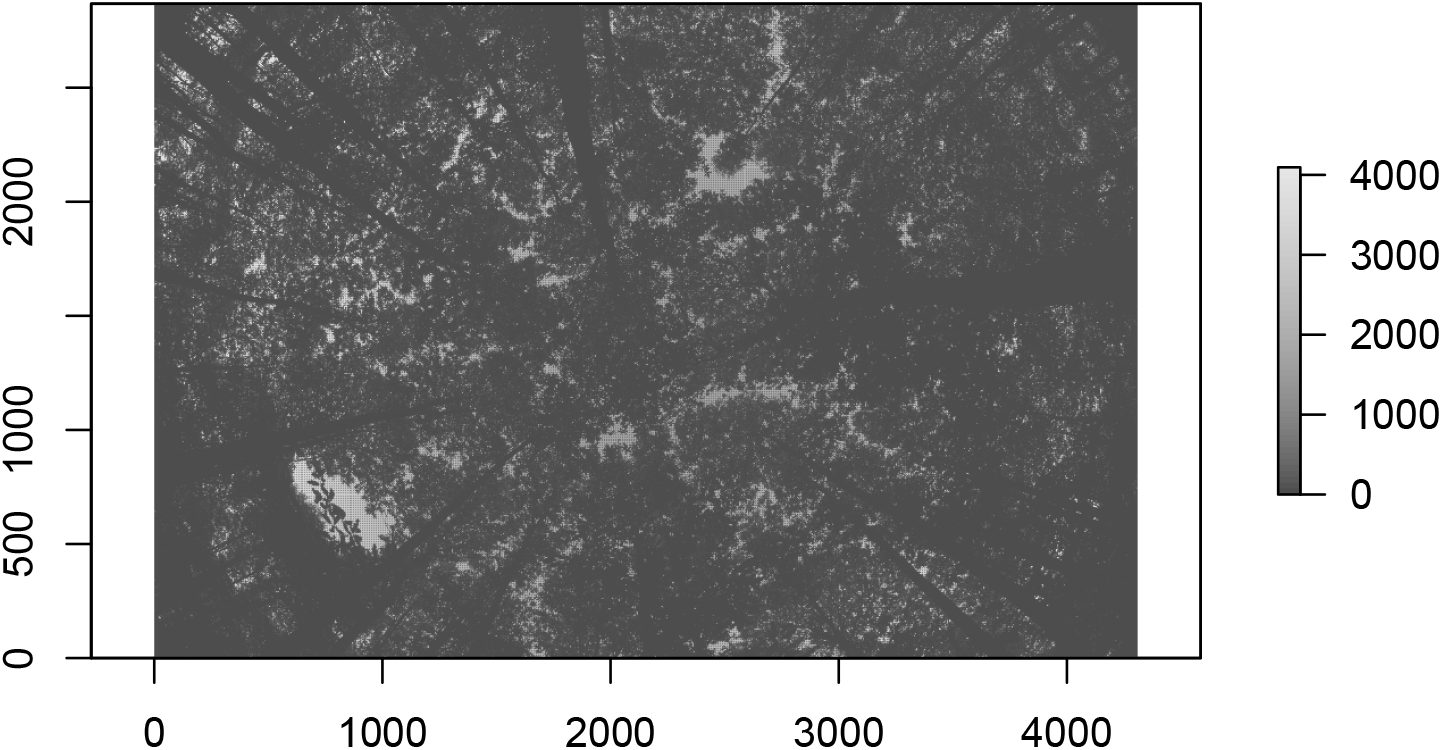

~~~
#> [1] ““
#> [2] “Filename: /Users/francescochianucci/Library/R/x86_64/4.1/library/bRaw/extdata/531539_AGŨ12_531
#> [3] “Timestamp: Fri Aug 24 19:26:10 2012”
#> [4] “Camera: Nikon D90”
#> [5] “ISŨ speed: 400”
#> [6] “Shutter: 1/60.0 sec”
#> [7] “Aperture: f/8.0”
#> [8] “Focal length: 10.5 mm”
#> [9] “Embedded ICC profile: no”
#> [10] “Number of raw images: 1”
#> [11] “Thumb size: 4288 x 2848”
#> [12] “Full size: 4352 x 2868”
#> [13] “Image size: 4310 x 2868”
#> [14] “Output size: 2868 x 4310”
#> [15] “Raw colors: 3”
#> [16] “Filter pattern: GB/RG”
#> [17] “Daylight multipliers: 2.008508 0.925194 1.076160”
#> [18] “Camera multipliers: 554.000000 256.000000 286.000000 256.000000”
~~~

When zooming-in the image it is possible to see the demosaiced raw pattern:

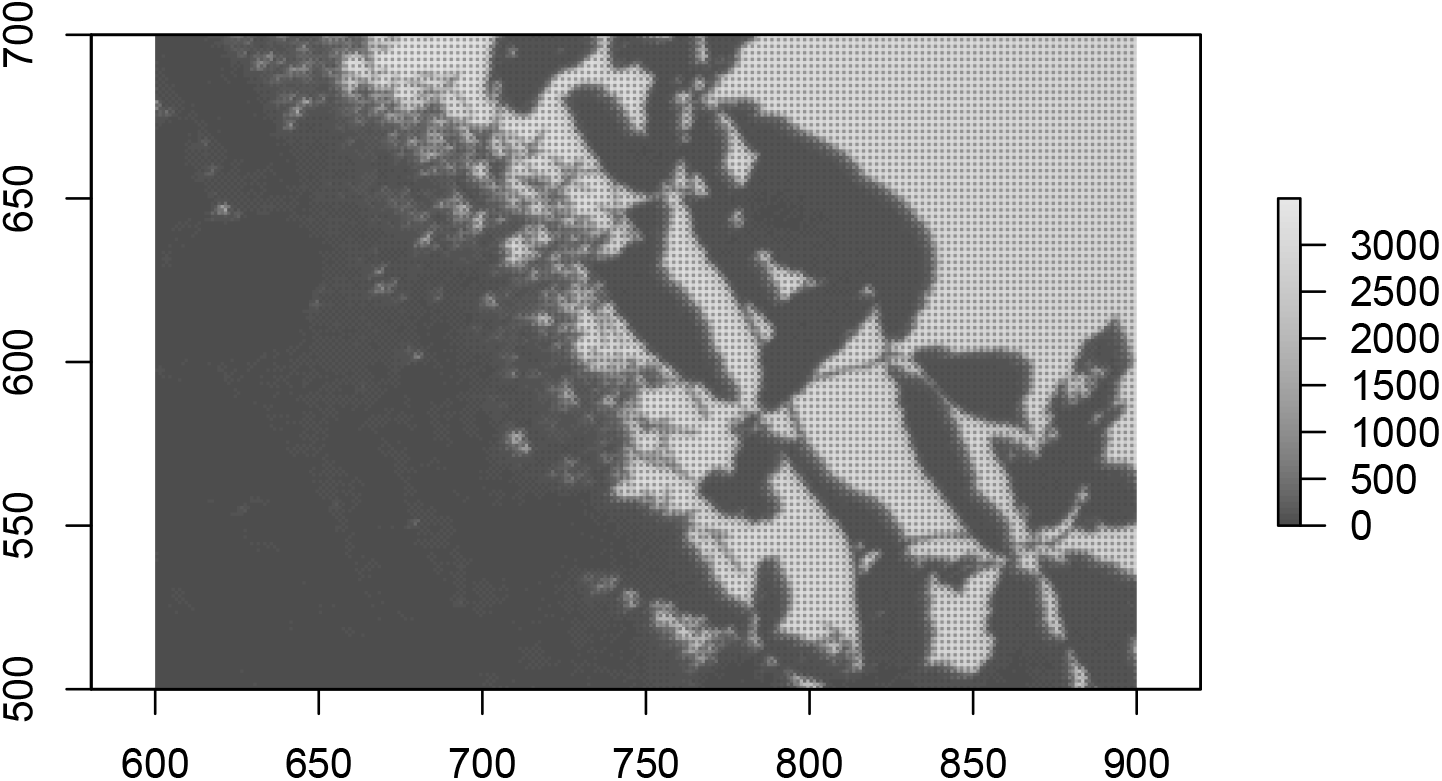

Using the function `raw_blue()` it yields:

~~~
raw_blue(filename)
~~~

The results is a single channel, 8-bit JPEG image, with increased contrast (high dynamic range):

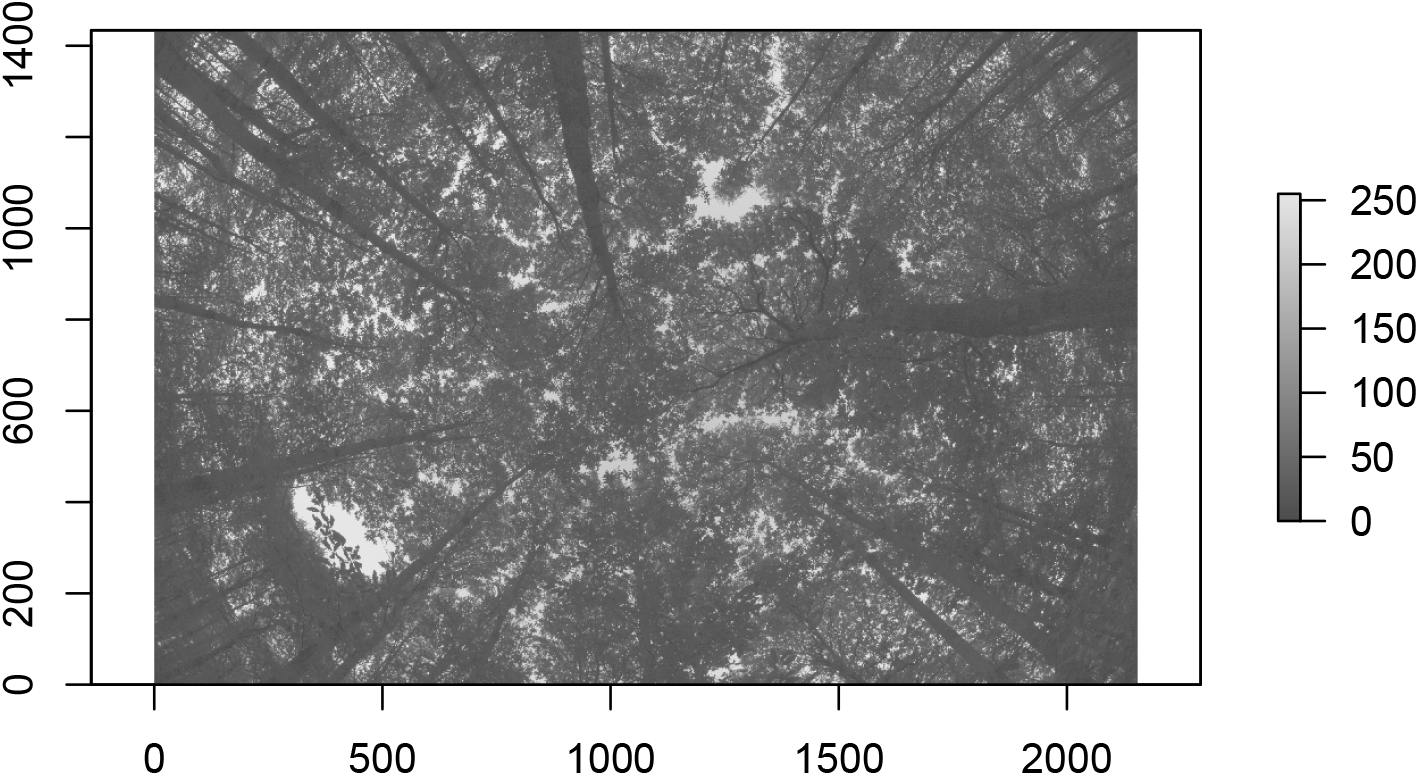

Looking at the histogram:

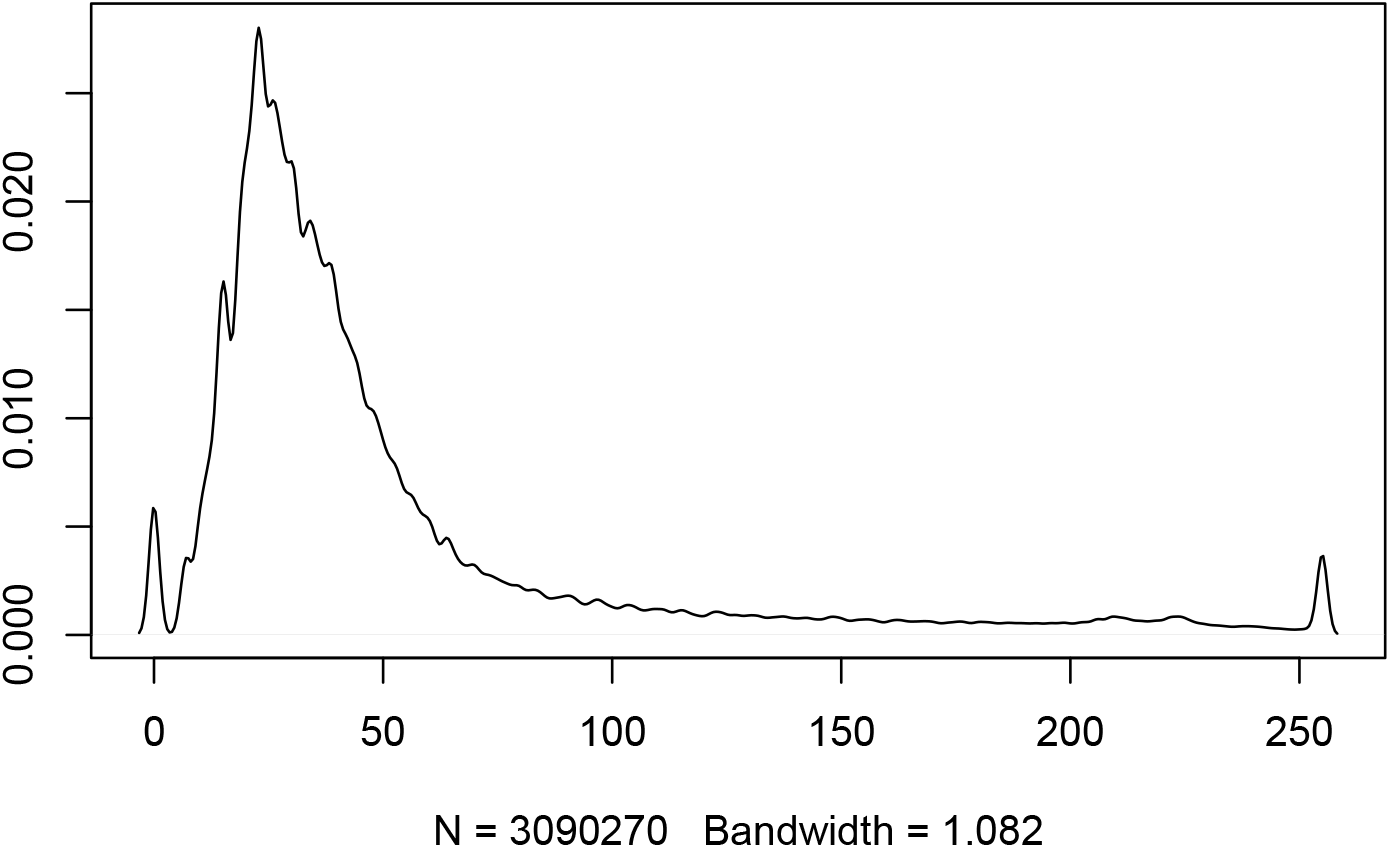

Here another example with a circular fisheye image, which needs setting a circular mask:

~~~
raw_blue(circ.img,circ.mask=list(xc=2144,yc=1424,rc=1050))
~~~

which yields:

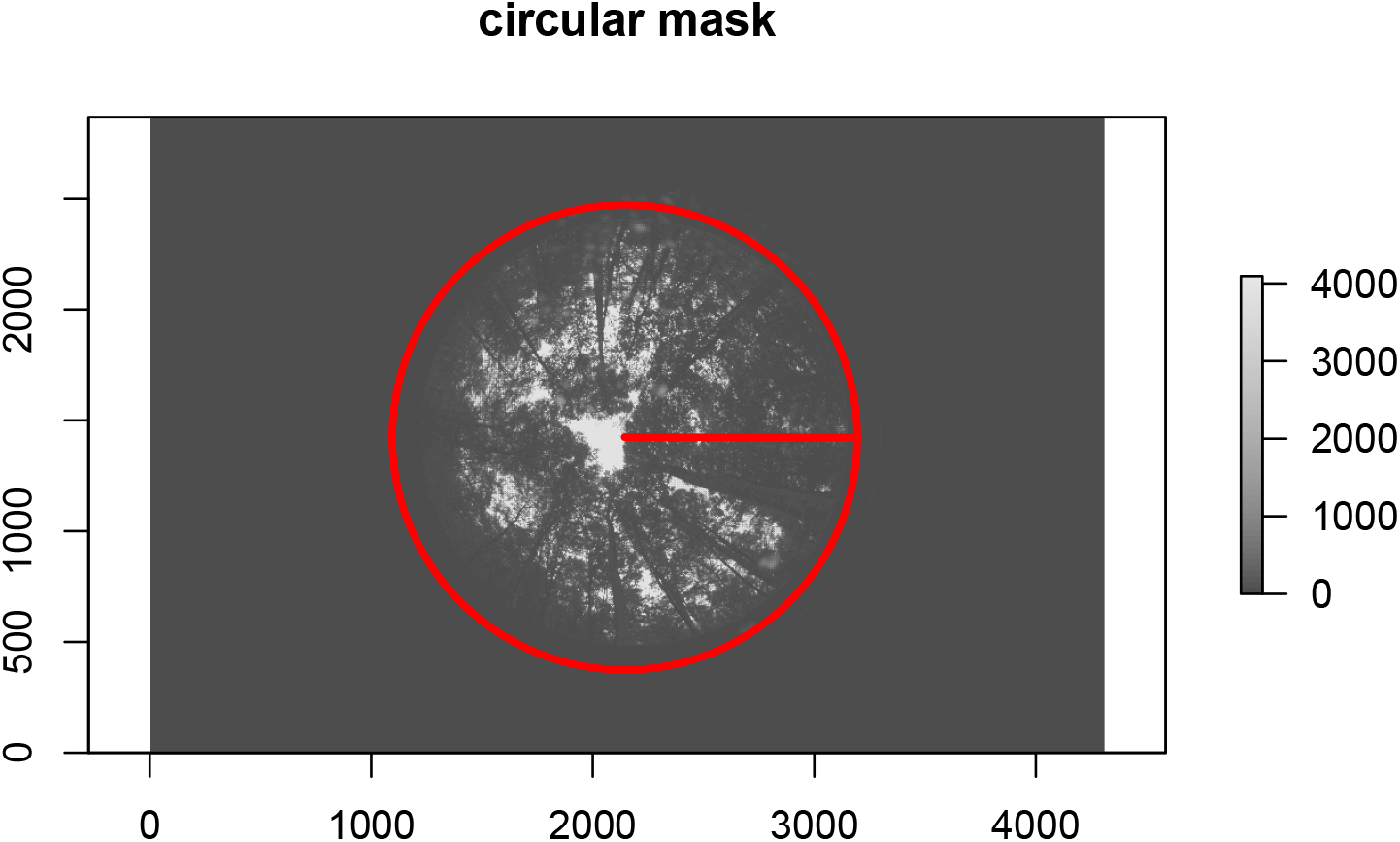

### Way forward

Once 8-bit JPEG images are derived from raw imagery, fisheye (either circular, full-frame) or 57° images can be processed using the hemispheR package (Chianucci and Macek (2022)). Cover photography images can be insted processed using the coveR package (Chianucci et al. (2022))

